# Mobility and infectiousness in the spatial spread of an emerging fungal pathogen

**DOI:** 10.1101/2020.05.07.082651

**Authors:** Kate E. Langwig, J. Paul White, Katy L. Parise, Heather M. Kaarakka, Jennifer A. Redell, John E. DePue, William H. Scullon, Jeffrey T. Foster, A. Marm Kilpatrick, Joseph R. Hoyt

**Affiliations:** Department of Biological Sciences, Virginia Polytechnic Institute, Blacksburg, VA, 24060 USA; Wisconsin Department of Natural Resources, Madison, WI, 53703 USA; Pathogen and Microbiome Institute, Northern Arizona University, Flagstaff, AZ 86011, USA; Michigan Department of Natural Resources, Baraga, MI 49870, USA; Michigan Department of Natural Resources, Norway, MI 49908, USA; Department of Ecology and Evolutionary Biology, University of California, Santa Cruz, CA 95064 USA

**Keywords:** migration, infectious disease, *Pseudogymnoascus destructans*, pathogen seasonality, white-nose syndrome

## Abstract

1. Emerging infectious diseases can have devastating effects on host communities, causing population collapse and species extinctions. The timing of novel pathogen arrival into naïve species communities can have consequential effects that shape the trajectory of epidemics through populations. Pathogen introductions are often presumed to occur when hosts are highly mobile. However, spread patterns can be influenced by a multitude of other factors including host body condition and infectiousness.
2. White-nose syndrome (WNS) is a seasonal emerging infectious disease of bats, which is caused by the fungal pathogen *Pseudogymnoascus destructans*. Within-site transmission of *P. destructans* primarily occurs over winter, however the influence of bat mobility and infectiousness on the seasonal timing of pathogen spread to new populations is unknown. We combined data on host population dynamics and pathogen transmission from 22 bat communities to investigate the timing of pathogen arrival and the consequences of varying pathogen arrival times on disease impacts.
3. We found that midwinter arrival of the fungus predominated spread patterns, suggesting that bats were most likely to spread *P. destructans* when they are highly infectious, but have reduced mobility. In communities where *P. destructans* was detected in early winter, one species suffered higher fungal burdens and experienced more severe declines than at sites where the pathogen was detected later in the winter, suggesting that the timing of pathogen introduction had consequential effects for some bat communities. We also found evidence of source-sink population dynamics over winter, suggesting some movement among sites occurs during hibernation, even though bats at northern latitudes were thought to be fairly immobile during this period. Winter emergence behavior symptomatic of white-nose syndrome may further exacerbate these winter bat movements to uninfected areas.
4. Our results suggest that low infectiousness during host migration may have reduced the rate of expansion of this deadly pathogen, and that elevated infectiousness during winter plays a key role in seasonal transmission. Furthermore, our results highlight the importance of both accurate estimation of the timing of pathogen spread and the consequences of varying arrival times to prevent and mitigate the effects of infectious diseases.

## 1. Introduction

The seasonality of pathogen spread is important for understanding, predicting, and controlling disease outbreaks. Pathogens infecting highly mobile hosts often have rapid rates of spread (Conner & Miller 2004; Altizer *et al*. 2006; Altizer, Bartel & Han 2011; Dalziel, Pourbohloul & Ellner 2013). However, key tradeoffs exist between mobility and disease, which can affect the likelihood that hosts spread pathogens during periods of high mobility (Kiesecker *et al*. 1999; Norris & Evans 2000; Wendland *et al*. 2010; Shakhar & Shakhar 2015). For example, migration can affect host body condition and immune status which could increase host susceptibility to infection (Altizer, Bartel & Han 2011). However, once a highly mobile host becomes infected, behavioral responses, such as sickness behavior, are likely to decrease mobility (Van Gils *et al*. 2007; Bouwman & Hawley 2010). Pathogen spread should therefore be most likely to occur when hosts are both highly infectious and mobile, but given tradeoffs caused by sickness behavior, peak mobility and infectiousness may not temporally align (Shakhar & Shakhar 2015). Understanding the relative importance of these factors in the seasonal timing of pathogen spread is critical because varying pathogen arrival times can have consequential effects on hosts that shape the trajectory of epidemics (Dalziel *et al*. 2018).

White-nose syndrome (WNS) is an annual seasonal epidemic that occurs in bat communities during winter (Langwig *et al*. 2015a; Hoyt *et al*. 2020a). The fungus that causes WNS, *Pseudogymnoascus destructans*, is psychrophilic and grows only at cool temperatures (1–20°C) (Verant *et al*. 2012) which restricts fungal replication and infection into bats’ epidermal tissue to periods when bats are in torpor (e.g. cool their body temperatures <20°C) (Meteyer *et al*. 2009; Langwig *et al*. 2015a; Langwig *et al*. 2016). White-nose syndrome disrupts bat physiology during hibernation, causing increased arousals that result in bats prematurely burning through fat stores before the end of winter (Lorch *et al*. 2011; Reeder *et al*. 2012; Warnecke *et al*. 2012; Warnecke *et al*. 2013; Verant *et al*. 2014). The spread of *P. destructans* has resulted in dramatic declines in bat communities across North America, and threatens several species with extinction (Frick *et al*. 2010; Langwig *et al*. 2012; Frick *et al*. 2015; Langwig *et al*. 2016). Seasonal changes in host behavior and physiology paired with seasonal differences in pathogen growth may influence the timing of pathogen spread to new communities.

The seasonal timing of *P. destructans* spread among sites is unknown but may occur during autumn when bats are highly mobile during mating and move among caves and mines (Davis & Hitchcock 1965; Cope & Humphrey 1977; Thomas, Fenton & Barclay 1979; Glover & Altringham 2008), which are known reservoirs of the fungus (Hoyt *et al*. 2020a). Autumn swarm behavior is an important source of gene flow when individuals move large distances between otherwise disconnected populations and contact among mating individuals could lead to pathogen transmission (Veith *et al*. 2004; Arnold 2007; Glover & Altringham 2008; Rossiter *et al*. 2012; Wilder, Kunz & Sorenson 2015). However, the spread of WNS has occurred relatively slowly compared with other disease systems with similarly mobile hosts. For example, WNS and West Nile virus were both introduced to New York, USA, but in 5 years West Nile virus had reached California whereas WNS spanned less than half of eastern North America (Kilpatrick *et al*. 2006; Kilpatrick, LaDeau & Marra 2007). In addition, *P. destructans* is less prevalent on bats during autumn, and infected individuals typically have reduced infection levels (i.e. 100–1000-fold lower than in midwinter; (Langwig *et al*. 2015a), Figure 1), suggesting that spread may occur outside of the highly mobile autumn period. If infectiousness is more important than mobility in *P. destructans* spread among sites, then pathogen introduction to new communities could occur during winter when fungal loads on bats are highest (Langwig *et al*. 2015a). However, most species are thought to be highly philopatric to sites and relatively sedentary during winter, particularly at northern latitudes where temperatures fall well below freezing (Davis & Hitchcock 1965; Davis 1970; Fenton & Barclay 1980; Fujita & Kunz 1984; Caceres & Barclay 2000). Therefore, further investigation is needed to disentangle the importance of host mobility and infectiousness in the spread of *P. destructans*.

**Figure 1.**
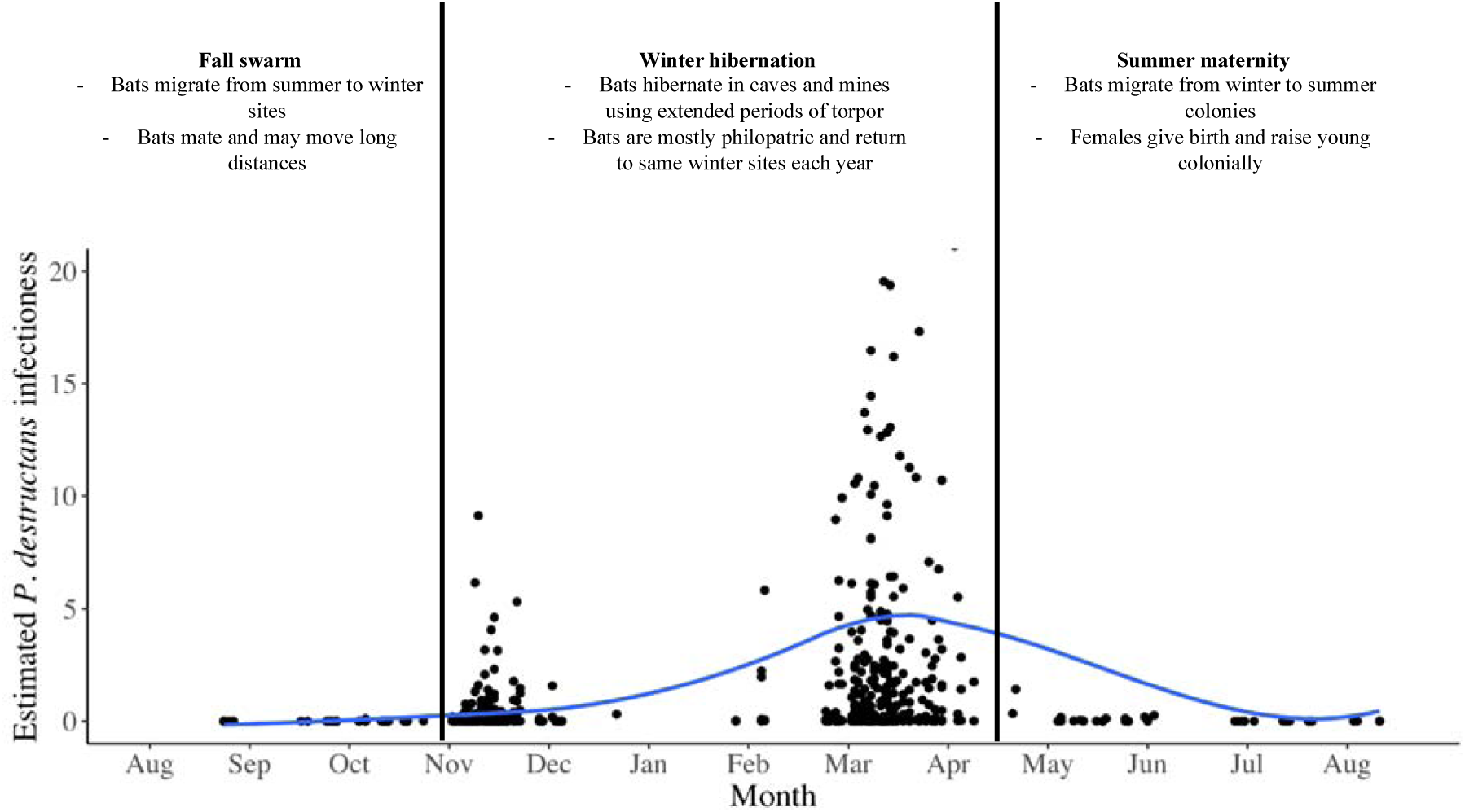
Mobility, migration, and infectiousness of temperate bats infected with *P. destructans*. Data from (Langwig *et al*. 2015a) and (Hoyt *et al*. 2020b) from little brown, northern long-eared, big brown, and tri-colored bats. Each estimate of infectiousness is derived from the mean fungal loads of all positive individuals of a given species captured on a given date (‘average fungal loads’) multiplied by the fraction of individuals of that species positive for *P. destructans* (‘prevalence’), multiplied by the proportional abundance of the species in the sample. This estimate is commonly referred to as ‘propagule pressure’ in invasion ecology (Lockwood, Cassey & Blackburn 2005). For example, northern long-eared bats, which are rare (typically comprising <5% of captures), but have high fungal loads and high prevalence would receive a lower estimate of infectiousness than little brown bats with equally high loads and prevalence but more commonly captured. Blue lines show loess curve smoothing to data points to ease visualization. Temperate hibernating bats are thought to have high mobility during fall swarm due to mating behavior and migration between summer and winter grounds while remaining more sedentary during winter while they hibernate (Davis & Hitchcock 1965; Thomas, Fenton & Barclay 1979; Thomas, Dorais & Bergeron 1990).

We investigated the timing of arrival of *P. destructans* into 22 hibernating bat communities in the Midwestern U.S. We hypothesized that pathogen invasion would be more likely to occur during winter when bats are less mobile but highly infectious, resulting in the first detections of *P. destructans* at each site occurring in late, rather than early winter. Given the potential importance of seasonally varying arrival times on disease dynamics, we also assessed the influence of timing of *P. destructans* introductions on disease severity, hypothesizing that earlier introductions of *P. destructans* would result in higher infection prevalence, fungal loads, and population impacts.

## 2. Methods

### 2.1 Study sites and data collection

We studied patterns of *P. destructans* arrival at 22 sites in the Midwestern U.S. Over a 5-year period, we visited each site twice per winter and collected data on population dynamics and infection status of four hibernating bat species (*Myotis lucifugus, Myotis septentrionalis, Eptesicus fuscus*, and *Perimyotis subflavus*). We used epidermal swab sampling to determine the presence and abundance of *P. destructans* on bats at two time points during hibernation (November - early hibernation, and March - late hibernation) in each year. During each visit, we counted all bats present and identified bats to the species level. In addition, we installed HOBO U23 Pro v2 temperature (+/- 0.2 C accuracy) and humidity (+/- 3.5 - 5%) loggers at 1–4 locations within a site to determine roost temperature and humidity.

### 2.2 Sample testing

We sampled bats using a standardized protocol (Langwig *et al*. 2015a) and stored swabs in RNAlater® for sample preservation until extraction. We tested samples for *P. destructans* DNA using real-time PCR (Muller *et al*. 2013) and quantified fungal loads based on the cycle threshold (C_t_) value to estimate a fungal load on each bat, with a cut-off of 40 cycles. Quantification of serial dilutions of the DNA from 10 ng to 1000 fg resulted in C_t_ scores ranging from 17.33 to 30.74 and a quantification relationship of C_t_ = -3.348* log_10_ (*P. destructans*[ng]) + 22.049, r^2^ = 0.986. We calculated prevalence as the proportion of bats of each species testing positive for *P. destructans* out of the number of individuals of that species sampled.

### 2.3 Statistical analysis

We used modified binomial power analyses to assess our ability to detect *P. destructans* arrival at each site where no positive samples were detected. We first calculated an expected early prevalence at each site where *P. destructans* was not detected in early hibernation based on a weighted mean prevalence. The weighted mean prevalence was calculated as the average prevalence of each species at sites where *P. destructans* was detected in early hibernation multiplied by the proportional abundance of a given species at each site (Table S1). We then calculated the probability of missing *P. destructans* at a site as the probability of getting all negatives in bats given the expected prevalence at a site multiplied by the probability of missing *P. destructans* in the unsampled bats (calculation shown in Appendix).

We investigated the effect of timing of *P. destructans* arrival on late winter prevalence by fitting a generalized linear mixed model with a binomial distribution and a logit link. We included fixed effects for the timing of *P. destructans* detection (early or late hibernation) interacting with the effect of species, and included site as a random effect. We also examined the effect of timing of *P. destructans* detection on fungal loads using a linear mixed model with species and timing of detection as interacting fixed effects, and site as a random effect. Lastly, we assessed the effect of the timing of *P. destructans* introduction on log_10_ bat population growth rates, calculated as the annual change in late winter counts at each site, using linear mixed models with species interacting with timing of detection as fixed effects and site as a random effect. We explored the inclusion of autumn prevalence as a continuous variable (0 for all sites and species where *P. destructans* was not detected, prevalence in each species ranged between 0 and 0.25 for sites where *P. destructans* was detected during autumn) and the results were qualitatively similar to those shown, so we present the discretized (early or late winter *P. destructans* detection) for simplicity of visualization and interpretation. Lastly, we investigated evidence of overwinter movements of bats among sites. At each site, we calculated our response variable, overwinter *λ*, as the proportional change in the number of bats of each species at a site by dividing the late hibernation count by the early hibernation count. We then used a linear mixed model to assess the relationship between log_10_(overwinter *λ*) and log_10_(early hibernation colony size) with an additive or interactive effect of species and site as a random effect.

## 3. Results

During early hibernation, we sampled 706 total bats from 22 sites in the year that *P. destructans* was first detected (site mean +/- SE = 67.4 +/- 1.79)(Table S1). We first detected *P. destructans* in early hibernation at 32% of sites (7/22), whereas in 68% of sites (15/22), the detection of *P. destructans* first occurred during our late hibernation visit (Figure 2). A total of 468 bats (site mean +/- SE = 31.1 +/- 1.25) were sampled from sites that were negative for *P. destructans* during early hibernation and became positive later in the same winter. After accounting for the relatively high proportion of bats sampled in each site (an average of 38% of the total bats at each site) and variable estimated “true” prevalence among sites (weighted mean prevalence of the species sampled from sites where *P. destructans* was detected in early hibernation: mean = 0.07, range = 0.05, 0.10), the mean probability of missing *P. destructans* at each negative site was 0.063 (median = 0.039, range: 0.005–0.3266), suggesting fairly good power to detect *P. destructans* presence.

**Figure 2.**
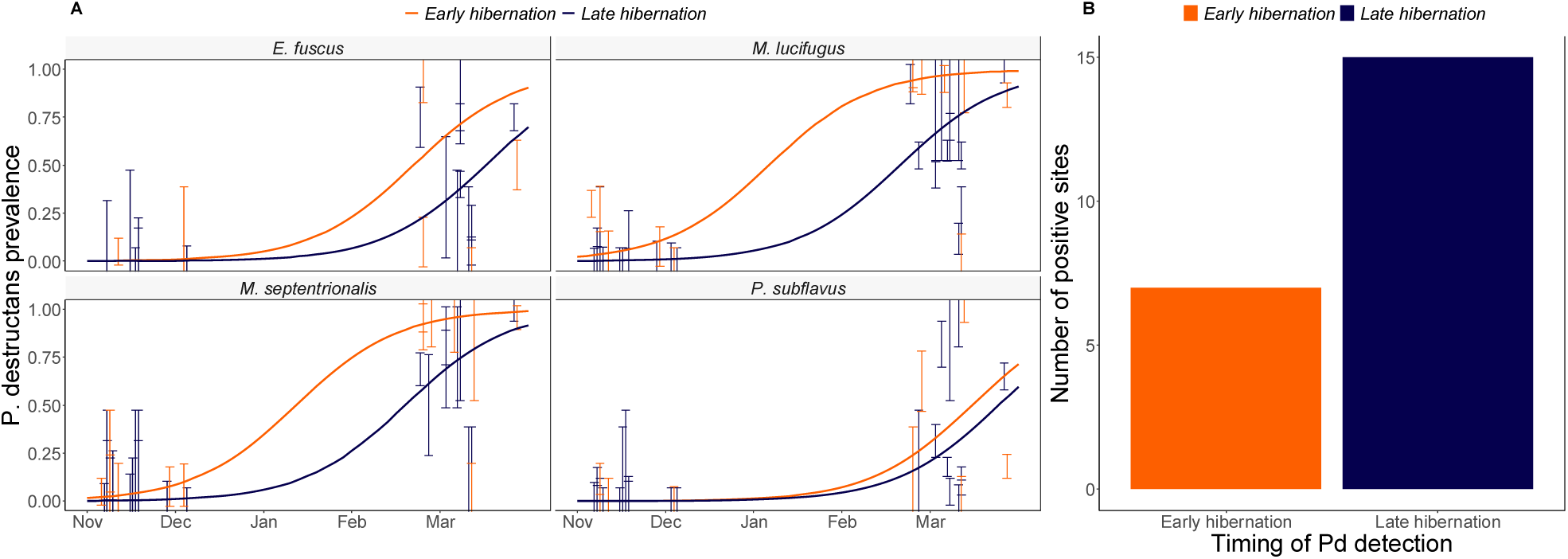
(a) Change in overwinter prevalence of *Pseudogymnoascus destructans* in the year that *P. destructans* was first detected at a site. Lines shows fits of a generalized linear mixed model with a binomial distribution with timing of first detection interacting with species and an additive effect of date with site as a random effect. (b) Twice as many sites had the first detection of *P. destructans* in late winter as in early winter.

Across sites where *P. destructans* was first detected during early hibernation, we were more likely to detect *P. destructans* on *M. lucifugus, M. septentrionalis*, and *E. fuscus* than on *P. subflavus* (*P. subflavus* intercept: -1.82 +/- 0.66, *M. lucifugus* coeff: 1.79 +/- 0.38 P < 0.0001, *E. fuscus* coeff: 1.40 +/- 0.51, P = 0.006, *M. septentrionalis* coef: 1.44 +/- 0.39, P = 0.0002, Appendix). In sites where *P. destructans* was first detected in late hibernation, we were also less likely to detect *P. destructans* on *P. subflavus* than on any other species (*P. subflavus* intercept: -1.69 +/- 0.31, *M. lucifugus* coeff: 0.94 +/- 0.22 P < 0.0001, *E. fuscus* coeff: 0.66 +/- 0.30, P = 0.0264, *M. septentrionalis* coeff: 0.84 +/- 0.26, P = 0.0014, Appendix).

The timing of *P. destructans* introduction influenced disease dynamics and population impacts for some, but not all, bat species (Figure 3). For *M. septentrionalis*, prevalence of *P. destructans*, fungal loads, and population impacts during late winter were higher at sites where *P. destructans* was first detected in early hibernation (one-tailed P-values = 0.032, 0.005, 0.0004, respectively, Appendix). For *M. lucifugus*, prevalence was also higher in sites where *P. destructans* was first detected in early hibernation (one-tailed P = 0.04, Appendix), although the effect on fungal loads and population impacts was less clear (Appendix). There was no clear effect of the timing of *P. destructans* introduction on prevalence, fungal loads, or population impacts for either *E. fuscus* or *P. subflavus*, possibly because these species had lower prevalence and fungal loads than *M. septentrionalis* and *M. lucifugus* in the first year of detection (Figure 3, Appendix).

**Figure 3.**
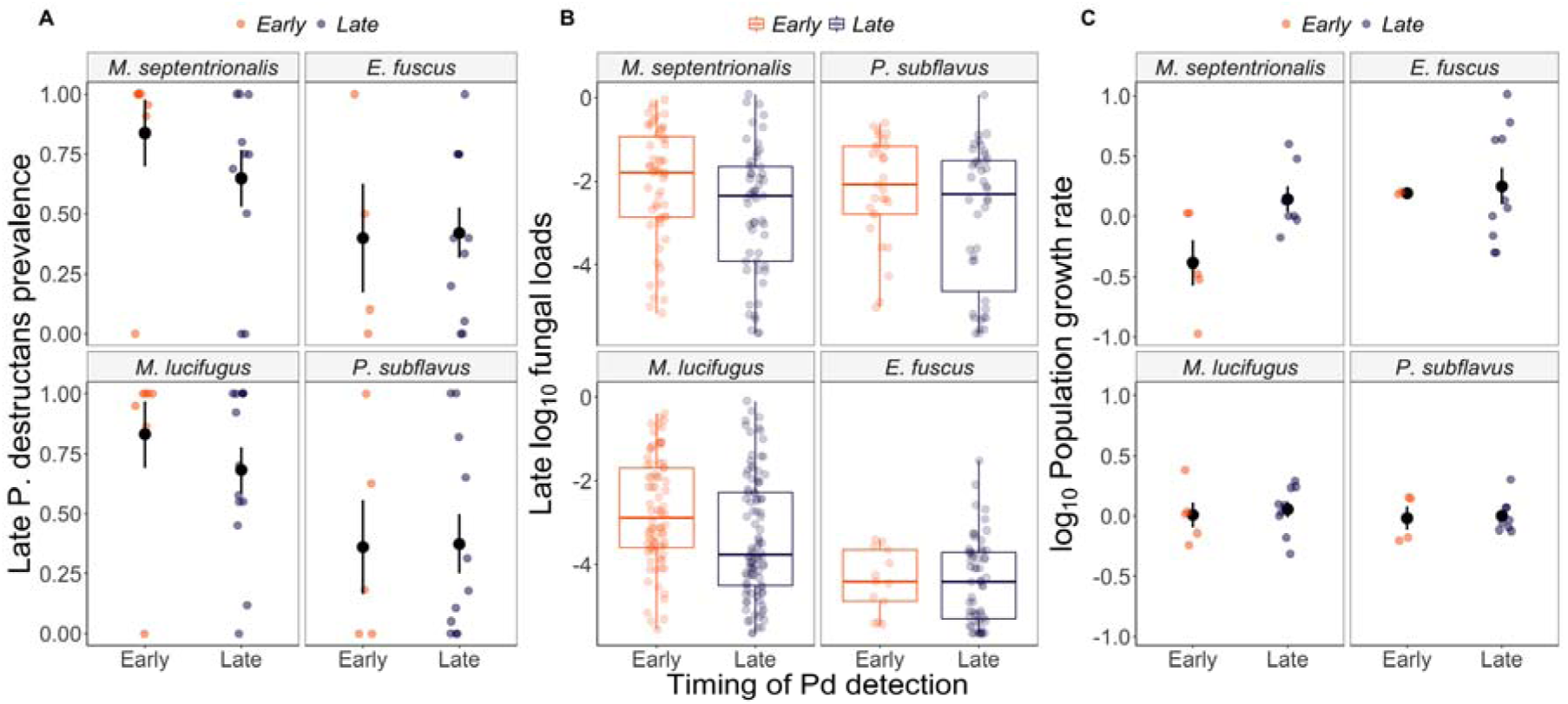
The timing of *Pseudogymnoascus destructans* introduction (early or late hibernation) influences late winter infection prevalence +/- SE (a), late winter fungal loads (b), and annual population impacts +/- SE (from late winter in the year that *P. destructans* arrives to the subsequent winter) (c), in some species. In sites where *P. destructans* was first detected during late hibernation *M. septentrionalis* had significantly higher infection prevalence, fungal loads, and population impacts by the end of winter in the first year of WNS.

We found that prior to the arrival of WNS, there were detectable changes in population counts between early and late hibernation (Figure 4). Across species, as colony size decreased in early hibernation, immigration increased (early hibernation colony size coeff: -0.15 +/- 0.06, t = -2.66) such that smaller colonies had proportionally more immigrants than larger colonies, indicative of a general spreading out of bats across sites during the hibernation season. We found no support for an interactive model over an additive model (ΔAIC = -9.63), suggesting no clear indication that the slope of this effect differed among species. The changes in overwinter counts occurred despite frigid minimum temperatures that were consistently below 5°C throughout this period (Figure S1), suggesting that some movement among hibernacula continues to occur during winter when bats are most infected with *P. destructans*.

**Figure 4.**
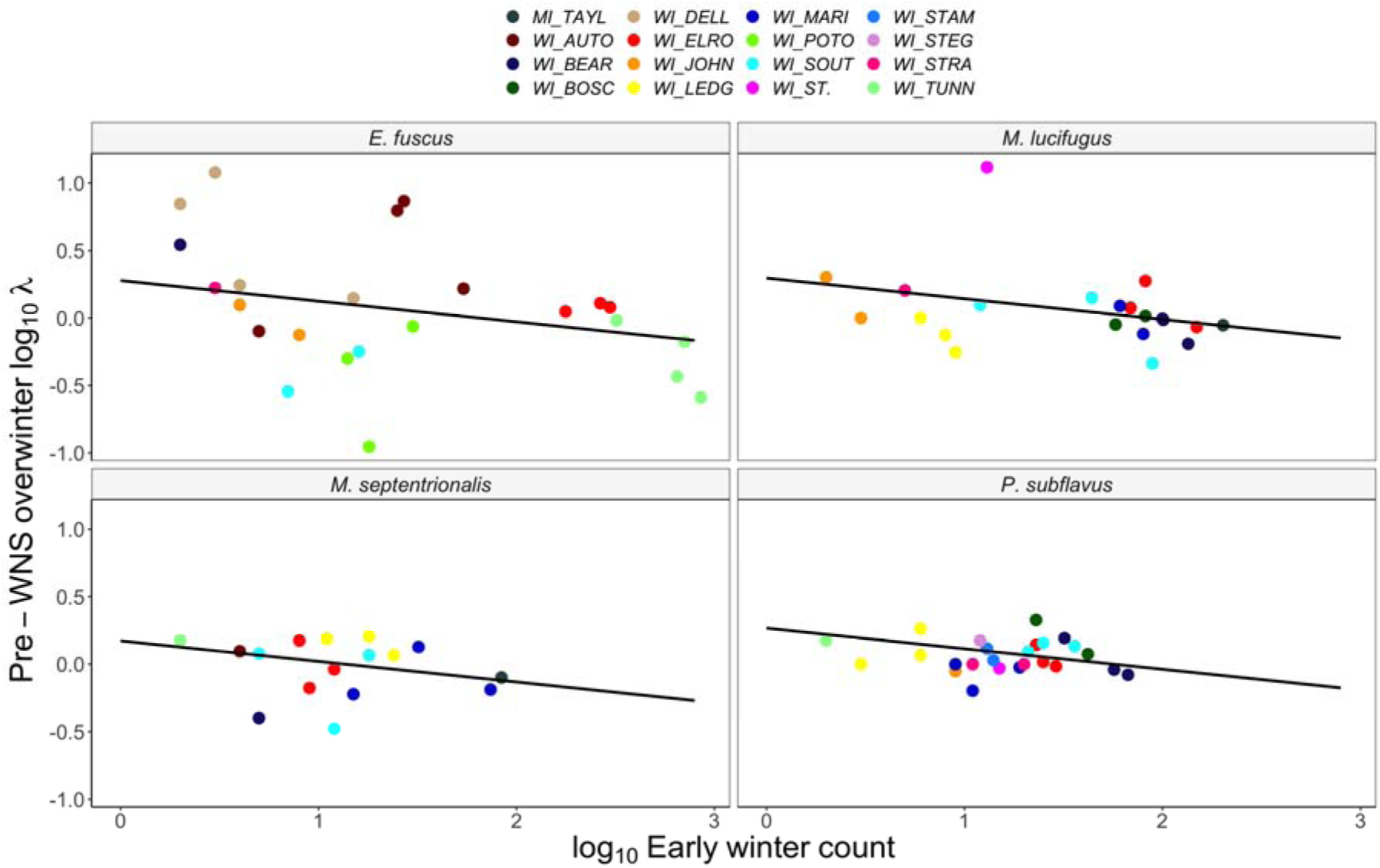
Pre-WNS immigration rates show dynamic populations during winter. Panels show bat species and colors indicate different bat populations. Lines show the fitted relationship between log_10_ early winter counts of bats and the rate of change in the same population over winter. Despite cold (<5°C) temperatures, there was considerable immigration and emigration among hibernation sites, consistent with bats moving from larger to smaller colonies over winter.

Lastly, we explored the effect of multiple covariates on the probability that *P. destructans* arrived at a site outside of or during the hibernation season, including abiotic (vapor pressure deficit and temperature) and biotic (abundance of certain species, species richness). We found no clear effects that any of these variables modified the probability that *P. destructans* was first detected at a site during early hibernation (Figure S2, Appendix).

## 4. Discussion

Our results indicate that host fungal burdens were important in determining the timing of pathogen spread. We found that *P. destructans* was more likely to be introduced to sites during winter hibernation than during autumn, suggesting that elevated infectiousness overwinter outweighed the increase in activity and long-distance movements that occur during autumn. Bats were previously thought to be restricted in their movement among sites during hibernation due to extremely cold temperatures (< - 5C; Fig S1) and a lack of prey availability. However, we found evidence that limited movement may be occurring among hibernacula during the winter, and given their increased levels fungal loads and prevalence, provide a higher probability of successful invasion to new sites. The lower fungal loads and prevalence on bats during autumn when they are highly mobile has likely decreased the rate of geographic expansion of the fungus as the majority of spread occurs during the hibernation period. As *P. destructans* moves into less connected western bat populations (Wilder, Kunz & Sorenson 2015; Lorch *et al*. 2016), spread could be further dampened by larger distances among colonies (Wilder, Kunz & Sorenson 2015; Weller *et al*. 2018).

Although the absolute absence of a pathogen is difficult to confirm, large sample sizes paired with the differential consequences of the timing of *P. destructans* introduction on disease dynamics, suggest that we did not simply fail to detect *P. destructans* during early winter. Our results also demonstrate that the timing of *P. destructans* introduction was more important for some species than others. Early introduction of *P. destructans* resulted in higher mortality in *M. septentrionalis* populations, but not *M. lucifugus*. Previous laboratory studies have shown that infection duration prior to death was 90-120 days for *M. lucifugus*. The absence of clear mortality in *M. lucifugus* populations while *M. septentrionalis* suffered from higher impacts suggests that *M. septentrionalis* may have a shorter time from infection until death. Infection in *P. subflavus* and *E. fuscus* were lower in the first year of WNS than *M. spp*., likely due to low environmental contamination (Langwig *et al*. 2015b; Hoyt *et al*. 2020a) and fewer contacts over the hibernation period (Hoyt *et al*. 2018).

We found evidence of some overwinter movement among hibernacula prior to the arrival of WNS, with a general trend of bats spreading out from larger to smaller colony size sites. Other studies have found that some species (especially the larger *E. fuscus*) can move in very cold temperatures (Klüg-Baerwald *et al*. 2016), although winter flight of *Myotis* spp. is thought to be a relatively rare occurrence (Davis & Hitchcock 1965). We cannot exclude the possibility that bats in larger sites became more difficult to detect in late winter while bats in smaller sites became more apparent. However, sites were selected for inclusion in this study for their relative simplicity in searching and ability to census all bats present (e.g. they did not have sections that we were unable to access and many were square tunnels with no obvious hiding places). As a result, we think that failed detections are unlikely to fully explain this relationship. In addition, higher overwinter mortality is unlikely to be a potential explanation as there is no evidence to suggest elevated mortality in sites with larger population sizes prior to the arrival of WNS (Langwig *et al*. 2012). Interestingly, symptoms of WNS include mid-hibernation emergence onto the winter landscape (Carr, Bernard & Stiver 2014; Bernard & McCracken 2017). This sickness behavior may increase movements between hibernacula and the spread of *P. destructans*.

We found no clear effects of any covariate in explaining differences in timing of arrival. However, we environmental conditions of hibernacula may be important determinants of *P. destructans* establishment into new communities outside of the hibernation season, as is suggested by the directionality and magnitude of a trend where warmer and wetter hibernacula tended to have a higher probability of autumn introduction. While bats with low fungal loads are likely moving among uninfected hibernacula during autumn, vapor pressure deficit and potentially temperature appear to be important in determining whether *P. destructans* can successfully establish in a population (Lilley, Anttila & Ruokolainen 2018). Given the limited number of sites where *P. destructans* arrived outside of the hibernation season, it was difficult to ascertain the importance of a number of factors that have been implicated by other studies (Wilder *et al*. 2011; Maher *et al*. 2012; Wilder, Kunz & Sorenson 2015). For example, species-specific differences in sociality and transmission (Hoyt *et al*. 2018) may be important in determining whether a lightly infected bat could introduce *P. destructans* during fall, although we found no clear effects of species abundance or composition on the timing of first detection of *P. destructans*. Additional research is needed to determine which species and site-specific differences may influence the timing of pathogen spread.

Importantly, many studies rely on disease stage (e.g. invading, epidemic or endemic) to draw inferences about geographic disease risk, population declines, and community persistence. Given the high probability of *P. destructans* introduction to new sites during winter, early and midwinter surveys could miss the introduction of *P. destructans*, and thus falsely conclude pathogen absence during the first year of arrival. Our results suggest that the arrival time of *P. destructans* can substantially influence dynamics, and therefore could be responsible for unexplained variance in transmission, impacts, and structure of remnant populations when sites are grouped by disease stage.

Our ability to predict the timing and patterns of pathogen spread are fundamental to the prevention and control of infectious disease outbreaks. The differential timing of initial pathogen arrival can have important effects on disease dynamics and lasting impacts to populations. This study suggests that host infectiousness is an important factor in determining successful pathogen spread and that incorrectly attributing pathogen spread to periods of high mobility would have masked the true underlying causes of among site variation in transmission and population impacts. Future studies examining the spatial and seasonal patterns of pathogen movement should consider the multitude of factors that might influence spread patterns.

## Acknowledgements

We thank Steffany Yamada, MI DNR, Eric McMaster and the many landowners for site access. The research was funded by NSF grants DEB-1115895, DEB-1336290, DEB-1911853 and the USFWS (F17AP00591). The authors have no competing interests to declare.

## Author Contributions

KEL analyzed the data and wrote the original draft; KEL, AMK and JRH revised the manuscript with input from all authors; KLP and JTF tested the samples and KEL, JPW, HMK, JAR, JED, WHS, AMK, and JRH collected the data.

## Data Availability Statement

Data will archived at VTData Repository upon manuscript acceptance or reviewer request.

## References

Altizer, S., Bartel, R. & Han, B.A. (2011) Animal Migration and Infectious Disease Risk. Science, 331, 296–302.

Altizer, S., Dobson, A., Hosseini, P., Hudson, P., Pascual, M. & Rohani, P. (2006) Seasonality and the dynamics of infectious diseases. Ecology Letters, 9, 467–484.

Arnold, B.D. (2007) Population structure and sex-biased dispersal in the forest dwelling vespertilionid bat, Myotis septentrionalis. The American Midland Naturalist, 157, 374–384.

Bernard, R.F. & McCracken, G.F. (2017) Winter behavior of bats and the progression of white-nose syndrome in the southeastern United States. Ecology and Evolution, 7, 1487–1496.

Bouwman, K.M. & Hawley, D.M. (2010) Sickness behaviour acting as an evolutionary trap? Male house finches preferentially feed near diseased conspecifics. Biology Letters, 6, 462–465.

Caceres, M.C. & Barclay, R.M.R. (2000) Myotis septentrionalis. Mammalian Species, 1–4.

Carr, J.A., Bernard, R.F. & Stiver, W.H. (2014) Unusual bat behavior during winter in Great Smoky Mountains National Park. Southeastern naturalist, 13, N18–N21.

Conner, M.M. & Miller, M.W. (2004) Movement patterns and spatial epidemiology of a prion disease in mule deer population units. Ecological Applications, 14, 1870–1881.

Cope, J.B. & Humphrey, S.R. (1977) SPRING AND AUTUMN SWARMING BEHAVIOR IN INDIANA BAT, MYOTIS-SODALIS. Journal of Mammalogy, 58, 93–95.

Dalziel, B.D., Kissler, S., Gog, J.R., Viboud, C., Bjørnstad, O.N., Metcalf, C.J.E. & Grenfell, B.T. (2018) Urbanization and humidity shape the intensity of influenza epidemics in US cities. Science, 362, 75–79.

Dalziel, B.D., Pourbohloul, B. & Ellner, S.P. (2013) Human mobility patterns predict divergent epidemic dynamics among cities. Proceedings of the Royal Society B: Biological Sciences, 280, 20130763.

Davis, W. (1970) Hibernation: ecology and physiological ecology. Biology of Bats, 1, 265 – 300.

Davis, W.H. & Hitchcock, H.B. (1965) Biology and migration of the bat, Myotis lucifugus, in New England. Journal of Mammalogy, 46, 296–313.

Fenton, M.B. & Barclay, R.M.R. (1980) Myotis lucifugus. Mammalian Species, 1–8.

Frick, W.F., Pollock, J.F., Hicks, A.C., Langwig, K.E., Reynolds, D.S., Turner, G.G., Butchkoski, C.M. & Kunz, T.H. (2010) An emerging disease causes regional population collapse of a common north american bat species. Science, 329, 679–682.

Frick, W.F., Puechmaille, S.J., Hoyt, J.R., Nickel, B.A., Langwig, K.E., Foster, J.T., Barlow, K.E., Bartonička, T., Feller, D., Haarsma, A.J., Herzog, C., Horacek, I., van der Kooij, J., Mulkens, B., Petrov, B., Reynolds, R., Rodrigues, L., Stihler, C.W., Turner, G.G. & Kilpatrick, A.M. (2015) Disease alters macroecological patterns of North American bats. Global Ecology and Biogeography, 24, 741–749.

Fujita, M.S. & Kunz, T.H. (1984) Pipistrellus subflavus. Mammalian Species, 1–6.

Glover, A.M. & Altringham, J.D. (2008) Cave selection and use by swarming bat species. Biological Conservation, 141, 1493–1504.

Hoyt, J.R., Langwig, K.E., Sun, K., Parise, K.L., Li, A., Wang, Y., Huang, X., Worledge, L., Miller, H. & White, J.P. (2020a) Environmental reservoir dynamics predict global infection patterns and population impacts for the fungal disease white-nose syndrome. Proceedings of the National Academy of Sciences, 117, 7255–7262.

Hoyt, J.R., Langwig, K.E., Sun, K., Parise, K.L., Li, A., Wang, Y., Huang, X., Worledge, L., Miller, H., White, J.P., Kaarakka, H.M., Redell, J.A., Görföl, T., Boldogh, S.A., Fukui, D., Sakuyama, M., Yachimori, S., Sato, A., Dalannast, M., Jargalsaikhan, A., Batbayar, N., Yovel, Y., Amichai, E., Natradze, I., Frick, W.F., Foster, J.T., Feng, J. & Kilpatrick, A.M. (2020b) Environmental reservoir dynamics predict global infection patterns and population impacts for the fungal disease white-nose syndrome. Proceedings of the National Academy of Sciences, 117, 7255.

Hoyt, J.R., Langwig, K.E., White, J.P., Kaarakka, H., Redell, J., Kurta, A., DePue, J., Scullon, W.H., Parise, K.L., Foster, J.T., Frick, W.F. & Kilpatrick, A.M. (2018) Cryptic connections illuminate pathogen transmission within community networks. Nature.

Kiesecker, J.M., Skelly, D.K., Beard, K.H. & Preisser, E. (1999) Behavioral reduction of infection risk. Proceedings of the National Academy of Sciences of the United States of America, 96, 9165–9168.

Kilpatrick, A.M., Chmura, A.A., Gibbons, D.W., Fleischer, R.C., Marra, P.P. & Daszak, P. (2006) Predicting the global spread of H5N1 avian influenza. Proceedings of the National Academy of Sciences of the United States of America, 103, 19368–19373.

Kilpatrick, A.M., LaDeau, S.L. & Marra, P.P. (2007) Ecology of west nile virus transmission and its impact on birds in the western hemisphere. Auk, 124, 1121–1136.

Klüg-Baerwald, B.J., Gower, L.E., Lausen, C. & Brigham, R. (2016) Environmental correlates and energetics of winter flight by bats in Southern Alberta, Canada. Canadian Journal of Zoology, 94, 829–836.

Langwig, K.E., Frick, W.F., Bried, J.T., Hicks, A.C., Kunz, T.H. & Marm Kilpatrick, A. (2012) Sociality, density-dependence and microclimates determine the persistence of populations suffering from a novel fungal disease, white-nose syndrome. Ecology Letters, 15, 1050–1057.

Langwig, K.E., Frick, W.F., Hoyt, J.R., Parise, K.L., Drees, K.P., Kunz, T.H., Foster, J.T. & Kilpatrick, A.M. (2016) Drivers of variation in species impacts for a multi-host fungal disease of bats. Phil. Trans. R. Soc. B, 371, 20150456.

Langwig, K.E., Frick, W.F., Reynolds, R., Parise, K.L., Drees, K.P., Hoyt, J.R., Cheng, T.L., Kunz, T.H., Foster, J.T. & Kilpatrick, A.M. (2015a) Host and pathogen ecology drive the seasonal dynamics of a fungal disease, white-nose syndrome. Proceedings of the Royal Society B: Biological Sciences, 282.

Langwig, K.E., Hoyt, J.R., Parise, K.L., Kath, J., Kirk, D., Frick, W.F., Foster, J.T. & Kilpatrick, A.M. (2015b) Invasion dynamics of white-nose syndrome white-nose syndrome fungus, midwestern United States, 2012-2014. Emerg Infect Dis, 21.

Lilley, T.M., Anttila, J. & Ruokolainen, L. (2018) Landscape structure and ecology influence the spread of a bat fungal disease. Functional Ecology, 32, 2483–2496.

Lockwood, J.L., Cassey, P. & Blackburn, T. (2005) The role of propagule pressure in explaining species invasions. Trends in Ecology & Evolution, 20, 223–228.

Lorch, J.M., Meteyer, C.U., Behr, M.J., Boyles, J.G., Cryan, P.M., Hicks, A.C., Ballmann, A.E., Coleman, J.T.H., Redell, D.N., Reeder, D.M. & Blehert, D.S. (2011) Experimental infection of bats with Geomyces destructans causes white-nose syndrome. Nature, 480, 376–378.

Lorch, J.M., Palmer, J.M., Lindner, D.L., Ballmann, A.E., George, K.G., Griffin, K., Knowles, S., Huckabee, J.R., Haman, K.H., Anderson, C.D., Becker, P.A., Buchanan, J.B., Foster, J.T. & Blehert, D.S. (2016) First Detection of Bat White-Nose Syndrome in Western North America. mSphere, 1, e00148–00116.

Maher, S.P., Kramer, A.M., Pulliam, J.T., Zokan, M.A., Bowden, S.E., Barton, H.D., Magori, K. & Drake, J.M. (2012) Spread of white-nose syndrome on a network regulated by geography and climate. Nature Communications, 3.

Meteyer, C.U., Buckles, E.L., Blehert, D.S., Hicks, A.C., Green, D.E., Shearn-Bochsler, V., Thomas, N.J., Gargas, A. & Behr, M.J. (2009) Histopathologic criteria to confirm white-nose syndrome in bats. Journal of Veterinary Diagnostic Investigation, 21, 411–414.

Muller, L.K., Lorch, J.M., Lindner, D.L., O’Connor, M., Gargas, A. & Blehert, D.S. (2013) Bat white-nose syndrome: a real-time TaqMan polymerase chain reaction test targeting the intergenic spacer region of Geomyces destructans. Mycologia, 105, 253–259.

Norris, K. & Evans, M.R. (2000) Ecological immunology: life history trade-offs and immune defense in birds. Behavioral Ecology, 11, 19–26.

Reeder, D.M., Frank, C.L., Turner, G.G., Meteyer, C.U., Kurta, A., Britzke, E.R., Vodzak, M.E., Darling, S.R., Stihler, C.W., Hicks, A.C., Jacob, R., Grieneisen, L.E., Brownlee, S.A., Muller, L.K. & Blehert, D.S. (2012) Frequent Arousal from Hibernation Linked to Severity of Infection and Mortality in Bats with White-Nose Syndrome. Plos One, 7.

Rossiter, S.J., Zubaid, A., MOHD-ADNAN, A., Struebig, M.J., Kunz, T.H., Gopal, S., Petit, E.J. & Kingston, T. (2012) Social organization and genetic structure: insights from codistributed bat populations. Molecular Ecology, 21, 647–661.

Shakhar, K. & Shakhar, G. (2015) Why Do We Feel Sick When Infected—Can Altruism Play a Role? Plos Biology, 13, e1002276.

Thomas, D., Dorais, M. & Bergeron, J. (1990) Winter energy budgets and cost of arousals for hibernating little brown bats, Myotis lucifugus. J Mammal, 71, 475 – 479.

Thomas, D.W., Fenton, M.B. & Barclay, R.M.R. (1979) Social behavior of the Little brown bat, Myotis lucifugus. I. mating behavior. Behavioral Ecology and Sociobiology, 6, 129–136.

Van Gils, J.A., Munster, V.J., Radersma, R., Liefhebber, D., Fouchier, R.A. & Klaassen, M. (2007) Hampered foraging and migratory performance in swans infected with low-pathogenic avian influenza A virus. Plos One, 2, e184.

Veith, M., Beer, N., Kiefer, A., Johannesen, J. & Seitz, A. (2004) The role of swarming sites for maintaining gene flow in the brown long-eared bat (Plecotus auritus). Heredity, 93, 342.

Verant, M.L., Boyles, J.G., Waldrep, W., Wibbelt, G. & Blehert, D.S. (2012) Temperature-dependent growth of Geomyces destructans, the fungus that causes bat white-nose syndrome. Plos One, 7.

Verant, M.L., Carol, M.U., Speakman, J.R., Cryan, P.M., Lorch, J.M. & Blehert, D.S. (2014) White-nose syndrome initiates a cascade of physiologic disturbances in the hibernating bat host. BMC physiology, 14, 10.

Warnecke, L., Turner, J.M., Bollinger, T.K., Lorch, J.M., Misra, V., Cryan, P.M., Wibbelt, G., Blehert, D.S. & Willis, C.K.R. (2012) Inoculation of bats with European Geomyces destructans supports the novel pathogen hypothesis for the origin of white-nose syndrome. Proceedings of the National Academy of Sciences of the United States of America, 109, 6999–7003.

Warnecke, L., Turner, J.M., Bollinger, T.K., Misra, V., Cryan, P.M., Blehert, D.S., Wibbelt, G. & Willis, C.K.R. (2013) Pathophysiology of white-nose syndrome in bats: a mechanistic model linking wing damage to mortality. Biology Letters, 9.

Weller, T.J., Rodhouse, T.J., Neubaum, D.J., Ormsbee, P.C., Dixon, R.D., Popp, D.L., Williams, J.A., Osborn, S.D., Rogers, B.W. & Beard, L.O. (2018) A review of bat hibernacula across the western United States: Implications for white-nose syndrome surveillance and management. Plos One, 13.

Wendland, L.D., Wooding, J., White, C.L., Demcovitz, D., Littell, R., Berish, J.D., Ozgul, A., Oli, M.K., Klein, P.A., Christman, M.C. & Brown, M.B. (2010) Social behavior drives the dynamics of respiratory disease in threatened tortoises. Ecology, 91, 1257–1262.

Wilder, A.P., Frick, W.F., Langwig, K.E. & Kunz, T.H. (2011) Risk factors associated with mortality from white-nose syndrome among hibernating bat colonies. Biology Letters, 7, 950–953.

Wilder, A.P., Kunz, T.H. & Sorenson, M.D. (2015) Population genetic structure of a common host predicts the spread of white-nose syndrome, an emerging infectious disease in bats. Molecular Ecology, 24, 5495–5506.

