## Supplement for "Mobility and infectiousness in the spatial spread of an emerging fungal pathogen"

Table S1. Characteristics of sites surveyed for *P. destructans*. The first two letters of the site designate the state where it is located. Sites were caves, mines, culverts, abandoned railroad tunnels, cellars, or excavated caves (caves that had with little or no natural entrances in modern times that were excavated by humans for recreation.) Species abbreviations are the first two letters of genus and the first two letters of species name. Year of detection is that winter year that *P. destructans* was first detected in the site. For example, winter 2015-2016 is labeled as 2016. Timing of detection is the sampling trip when *P. destructans* was first detected with Nov (November) as early hibernation and Mar (March) as late hibernation.

| **Site** | **Site type** | **Species Present** | **Year of detection** | **Timing of detection** | **Total population (Nov)** | **Proportion sampled (Nov)** |
| --- | --- | --- | --- | --- | --- | --- |
| WI_AUTO | cave | EPFU, MYLU | 2017 | Mar | 2 | 0.5 |
| WI_BEAR | cave | EPFU, MYLU, MYSE, PESU | 2016 | Mar | 179 | 0.240 |
| WI_BOSC | cave | MYLU MYSE, PESU | 2015 | Mar | 136 | 0.265 |
| WI_DELL | culvert | EPFU | 2017 | Mar | 0 | NA |
| MI_GLEN | mine | EPFU, MYLU, MYSE | 2015 | Mar | 315 | 0.083 |
| WI_HORS | cave | EPFU, MYLU, MYSE, PESU | 2015 | Mar | 1055 | 0.026 |
| WI_JOHN | basement | EPFU, MYLU, MYSE, PESU | 2015 | Mar | 21 | 0.952 |
| WI_MARI | excavated cave | MYLU MYSE, PESU | 2016 | Mar | 182 | 0.274 |
| WI_POTO | cave | EPFU, MYLU, MYSE | 2016 | Mar | 0 | NA |
| WI_SOUT | excavated cave | EPFU, MYLU, MYSE, PESU | 2017 | Mar | 76 | 0.658 |
| WI_ST. | mined cave | EPFU, MYLU, MYSE, PESU | 2015 | Mar | 249 | 0.161 |
| WI_STEG | mine | MYLU MYSE, PESU | 2015 | Mar | 24 | 0.75 |
| WI_STRA | mined tunnel | EPFU, MYLU, MYSE, PESU | 2015 | Mar | 25 | 0.96 |
| WI_TUNN | tunnel | EPFU, MYSE, PESU | 2017 | Mar | 885 | 0.0249 |
| IL_ZIMM | mine | EPFU, MYLU, MYSE, PESU, MYSO | 2013 | Mar | 17319 | 0.005 |
| MI_BEAT | mine | EPFU, MYLU, MYSE | 2015 | Nov | 81 | 0.321 |
| IL_BLAC | mine | EPFU, MYLU, MYSE, PESU, MYSO | 2013 | Nov | 7234 | 0.007 |
| WI_ELRO | tunnel | EPFU, MYLU, MYSE, PESU | 2016 | Nov | 143 | 0.315 |
| WI_LEDG | excavated cave | MYLU MYSE, PESU | 2017 | Nov | 51 | 0.608 |
| MI_MASS | mine | EPFU, MYLU, MYSE | 2015 | Nov | 165 | 0.158 |
| WI_STAM | cave | MYLU MYSE, PESU | 2015 | Nov | 23 | 0.783 |
| MI_TAYL | mine | MYLU, MYSE | 2016 | Nov | 270 | 0.148 |

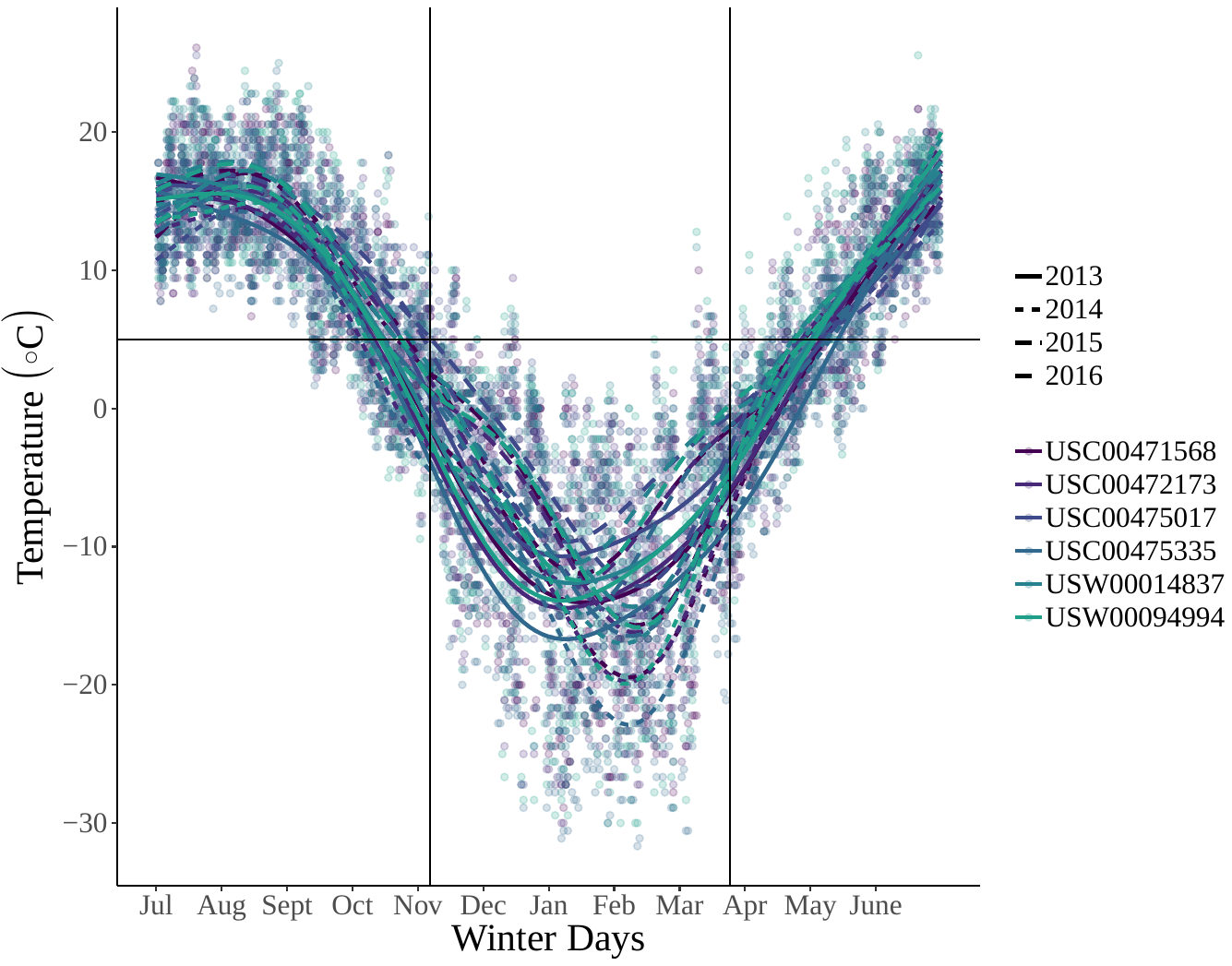

**Figure S1**

Minimum daily temperatures from weather stations collected within a 100-mile radius of hibernacula in the Midwestern U.S. The lines show generalized additive models fit to minimum temperatures at a weather station in a year and points show actual recorded minimum temperatures at that station. The solid horizontal lines show 5°C, and the vertical lines denote earliest and latest sampling and population census dates.

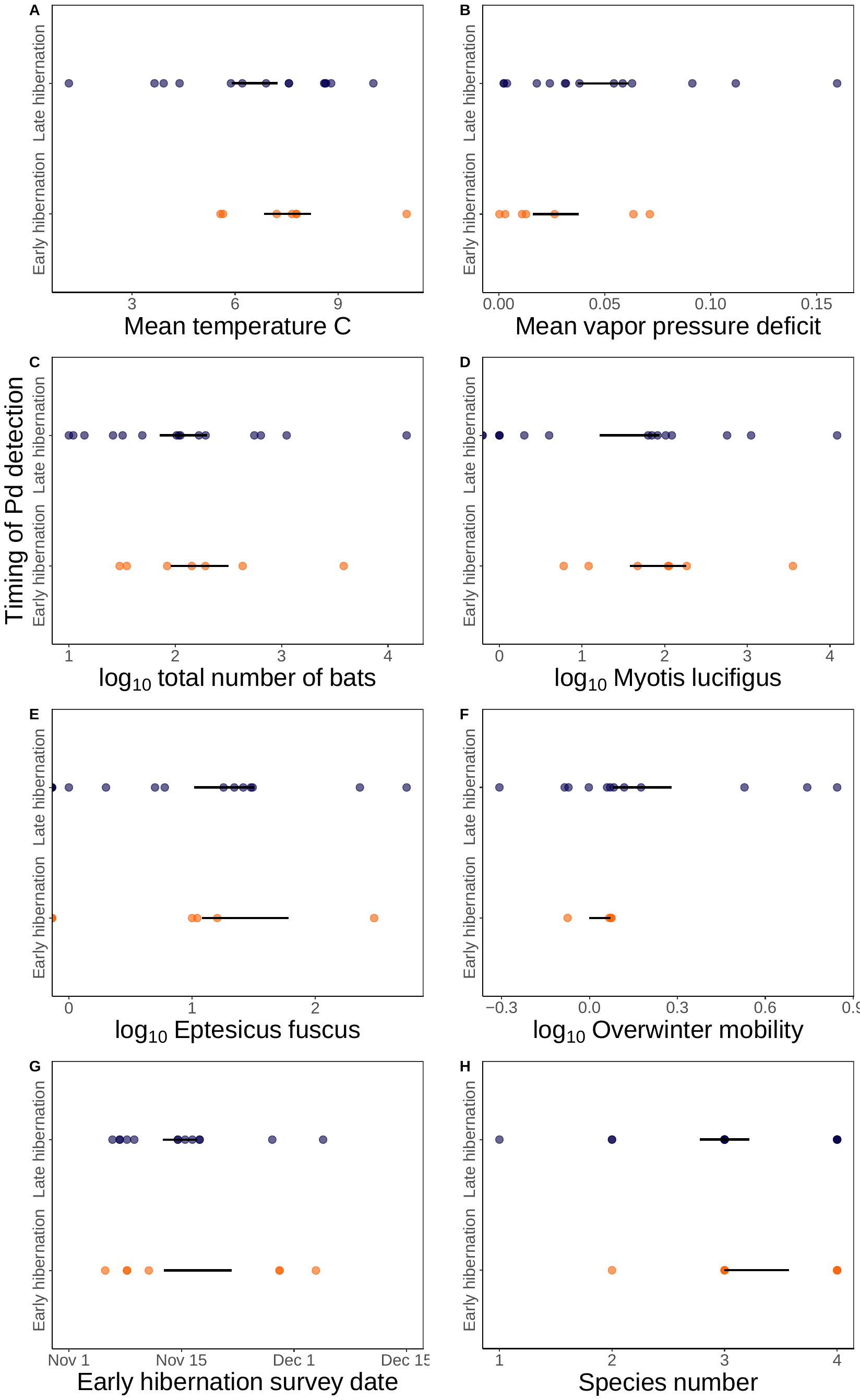

**Figure S2**

The effect of site covariates (a-h) on the timing of *Pseudogymnoascus destructans* detection across 22 hibernacula in the Midwestern U.S. No relationships were statistically significant (all p > 0.05), although mean temperature, mean vapor pressure deficit, and overwinter mobility (calculated as the average proportional change in counts pre-WNS) had large estimated coefficients such that sites where conditions were drier, of lower average temperature, and with higher overwinter mobility more likely to have *P. destructans* detected in late winter.
