## Appendix for "Mobility and infectiousness in the spatial spread of an emerging fungal pathogen"

R Notebook - Appendix for Langwig et al. Midwinter Arrival Ms


Code 

- Show All Code
- Hide All Code
- Download Rmd

### R Notebook - Appendix for Langwig et al. Midwinter Arrival Ms

- 1 POWER ANALYSES FOR DETECTION
  - 1.1 bats only calculations
    - 1.1.1 summarise detection probability across bats
- 2 WHICH SPECIES ARE WE MOST LIKELY TO DETECT PD ON?
  - 2.1 early winter analysis
  - 2.2 late winter analysis
- 3 CATEGORICAL ANALYSIS - HOW DOES DIFFERENTIAL ARRIVAL IMPACT LATE WINTER METRICS?
  - 3.1 prev in late winter
  - 3.2 loads in late winter
  - 3.3 lambda in late winter
- 4 CONTINUOUS ANALYSIS - SAME AS ABOVE BUT CONTINUOUS
  - 4.1 how are late winter loads affected by early prevalence?
  - 4.2 how is late winter prevalence affected by early prevalence?
  - 4.3 how are late winter impacts affected by early prevalence?
- 5 OVERWINTER MOBILITY
  - 5.1 Changes in counts over winter - site
  - 5.2 Does the log transformation of lambda help normality?
  - 5.3 Do model diagnostic plots look reasonable?
  - 5.4 If we preserve structure of data and use Gamma distribution do we get a similar finding?
- 6 NUMEROUS COVARIATES AND THE EFFECT ON TIMING OF DETECTION
  - 6.1 temperature
  - 6.2 vapor pressure deficit
  - 6.3 total bats
  - 6.4 number of MYLU
  - 6.5 number of EPFU
  - 6.6 overwinter mobility
  - 6.7 effect of early winter sampling date
  - 6.8 species richness

This is an R Markdown Notebook.

### 1 POWER ANALYSES FOR DETECTION

#### 1.1 bats only calculations


```
library(tidyverse)
library(dplyr)
data.site.summ = data %>%
  group_by(site,date)%>%
  mutate(tot.bats.sampled = sum(N, na.rm=T))%>%
  mutate(tot.bats.counted = sum(count, na.rm=T))%>%
  mutate(prop.sampled = tot.bats.sampled/tot.bats.counted)%>%
  mutate(weighted.average.prev.early = sum(weighted.average.early.num)/tot.bats.sampled)%>%
  mutate(site.prob.missing.traditional = dbinom(x=0,size=tot.bats.sampled,prob=weighted.average.prev.early))%>%
  mutate(site.prob.missing.all = (1 - weighted.average.prev.early)^(tot.bats.sampled)*(1-(1-weighted.average.prev.early)^(tot.bats.counted - tot.bats.sampled)))%>%
  filter(season=="hiber_earl" & t=="winter")
```


##### 1.1.1 summarise detection probability across bats


```
summary(missing.value.cals$site.prob.missing.all)
```


```
    Min.  1st Qu.   Median     Mean  3rd Qu.     Max. 
0.005171 0.017689 0.038582 0.063044 0.057758 0.326666
```

### 2 WHICH SPECIES ARE WE MOST LIKELY TO DETECT PD ON?

#### 2.1 early winter analysis


```
modS1=glmer(gd~species + (1|site) ,data=mw0.trim,family="binomial", subset=t=="fall");summary(modS1)
```


```
Generalized linear mixed model fit by maximum likelihood (Laplace Approximation) ['glmerMod']
 Family: binomial  ( logit )
Formula: gd ~ species + (1 | site)
   Data: mw0.trim
 Subset: t == "fall"

     AIC      BIC   logLik deviance df.resid 
   581.9    603.0   -286.0    571.9      496 

Scaled residuals: 
    Min      1Q  Median      3Q     Max 
-2.3486 -0.9002 -0.1708  0.9136  5.8562 

Random effects:
 Groups Name        Variance Std.Dev.
 site   (Intercept) 2.327    1.525   
Number of obs: 501, groups:  site, 7

Fixed effects:
            Estimate Std. Error z value Pr(>|z|)    
(Intercept)  -1.8225     0.6642  -2.744 0.006075 ** 
speciesMYLU   1.7925     0.3844   4.662 3.12e-06 ***
speciesEPFU   1.4013     0.5117   2.738 0.006175 ** 
speciesMYSE   1.4387     0.3909   3.681 0.000232 ***
---
Signif. codes:  0 ‘***’ 0.001 ‘**’ 0.01 ‘*’ 0.05 ‘.’ 0.1 ‘ ’ 1

Correlation of Fixed Effects:
            (Intr) spMYLU spEPFU
speciesMYLU -0.429              
speciesEPFU -0.351  0.627       
speciesMYSE -0.420  0.807  0.617
```

#### 2.2 late winter analysis


```
modS2=glmer(gd~species + (1|site) ,data=mw0.trim,family="binomial", subset=t=="winter");summary(modS2)
```


```
Generalized linear mixed model fit by maximum likelihood (Laplace Approximation) ['glmerMod']
 Family: binomial  ( logit )
Formula: gd ~ species + (1 | site)
   Data: mw0.trim
 Subset: t == "winter"

     AIC      BIC   logLik deviance df.resid 
  1032.6   1057.0   -511.3   1022.6      976 

Scaled residuals: 
    Min      1Q  Median      3Q     Max 
-1.3321 -0.6038 -0.3941  0.7507  4.6090 

Random effects:
 Groups Name        Variance Std.Dev.
 site   (Intercept) 0.8578   0.9262  
Number of obs: 981, groups:  site, 15

Fixed effects:
            Estimate Std. Error z value Pr(>|z|)    
(Intercept)  -1.6848     0.3090  -5.453 4.94e-08 ***
speciesMYLU   0.9344     0.2221   4.207 2.59e-05 ***
speciesEPFU   0.6558     0.2954   2.220  0.02641 *  
speciesMYSE   0.8375     0.2614   3.204  0.00135 ** 
---
Signif. codes:  0 ‘***’ 0.001 ‘**’ 0.01 ‘*’ 0.05 ‘.’ 0.1 ‘ ’ 1

Correlation of Fixed Effects:
            (Intr) spMYLU spEPFU
speciesMYLU -0.458              
speciesEPFU -0.458  0.470       
speciesMYSE -0.405  0.563  0.457
```

### 3 CATEGORICAL ANALYSIS - HOW DOES DIFFERENTIAL ARRIVAL IMPACT LATE WINTER METRICS?

#### 3.1 prev in late winter


```
mod2=glmer(gd~species*t + (1|site),weights=N, data=data.trim.late,family="binomial");summary(mod2)
```


```
Generalized linear mixed model fit by maximum likelihood (Laplace Approximation) ['glmerMod']
 Family: binomial  ( logit )
Formula: gd ~ species * t + (1 | site)
   Data: data.trim.late
Weights: N

     AIC      BIC   logLik deviance df.resid 
   258.9    278.9   -120.5    240.9       59 

Scaled residuals: 
    Min      1Q  Median      3Q     Max 
-3.3342 -0.4236  0.0902  0.5216  2.3033 

Random effects:
 Groups Name        Variance Std.Dev.
 site   (Intercept) 4.47     2.114   
Number of obs: 68, groups:  site, 22

Fixed effects:
                    Estimate Std. Error z value Pr(>|z|)    
(Intercept)           3.4459     1.0207   3.376 0.000735 ***
speciesEPFU          -3.2814     0.7489  -4.382 1.18e-05 ***
speciesMYLU          -0.4431     0.7344  -0.603 0.546278    
speciesPESU          -4.1628     0.7947  -5.238 1.62e-07 ***
twinter              -2.2458     1.2101  -1.856 0.063473 .  
speciesEPFU:twinter   1.5497     0.8804   1.760 0.078363 .  
speciesMYLU:twinter   0.2899     0.8339   0.348 0.728128    
speciesPESU:twinter   2.0029     0.9009   2.223 0.026205 *  
---
Signif. codes:  0 ‘***’ 0.001 ‘**’ 0.01 ‘*’ 0.05 ‘.’ 0.1 ‘ ’ 1

Correlation of Fixed Effects:
            (Intr) spEPFU spMYLU spPESU twintr sEPFU: sMYLU:
speciesEPFU -0.447                                          
speciesMYLU -0.453  0.588                                   
speciesPESU -0.482  0.590  0.563                            
twinter     -0.842  0.376  0.382  0.404                     
spcsEPFU:tw  0.377 -0.848 -0.499 -0.497 -0.444              
spcsMYLU:tw  0.400 -0.518 -0.881 -0.498 -0.449  0.587       
spcsPESU:tw  0.423 -0.519 -0.496 -0.878 -0.467  0.588  0.596
```


```
emmeans(mod2, pairwise ~ t|species)
```


```
$emmeans
species = MYSE:
 t      emmean    SE  df asymp.LCL asymp.UCL
 fall    3.446 1.021 Inf    1.4454     5.446
 winter  1.200 0.652 Inf   -0.0784     2.479

species = EPFU:
 t      emmean    SE  df asymp.LCL asymp.UCL
 fall    0.164 0.959 Inf   -1.7147     2.044
 winter -0.532 0.611 Inf   -1.7288     0.666

species = MYLU:
 t      emmean    SE  df asymp.LCL asymp.UCL
 fall    3.003 0.949 Inf    1.1419     4.864
 winter  1.047 0.597 Inf   -0.1239     2.218

species = PESU:
 t      emmean    SE  df asymp.LCL asymp.UCL
 fall   -0.717 0.944 Inf   -2.5678     1.134
 winter -0.960 0.606 Inf   -2.1477     0.228

Results are given on the logit (not the response) scale. 
Confidence level used: 0.95 

$contrasts
species = MYSE:
 contrast      estimate   SE  df z.ratio p.value
 fall - winter    2.246 1.21 Inf 1.856   0.0635 

species = EPFU:
 contrast      estimate   SE  df z.ratio p.value
 fall - winter    0.696 1.14 Inf 0.612   0.5404 

species = MYLU:
 contrast      estimate   SE  df z.ratio p.value
 fall - winter    1.956 1.12 Inf 1.746   0.0808 

species = PESU:
 contrast      estimate   SE  df z.ratio p.value
 fall - winter    0.243 1.12 Inf 0.217   0.8286 

Results are given on the log odds ratio (not the response) scale.
```

#### 3.2 loads in late winter


```
mod3=lmer(lgdL~species*t + (1|site), data=mw0.trim.late);summary(mod3)
```


```
Linear mixed model fit by REML ['lmerMod']
Formula: lgdL ~ species * t + (1 | site)
   Data: mw0.trim.late

REML criterion at convergence: 1348.1

Scaled residuals: 
     Min       1Q   Median       3Q      Max 
-2.90453 -0.62385  0.02865  0.58963  3.07815 

Random effects:
 Groups   Name        Variance Std.Dev.
 site     (Intercept) 1.021    1.011   
 Residual             1.194    1.093   
Number of obs: 429, groups:  site, 21

Fixed effects:
                    Estimate Std. Error t value
(Intercept)          -1.6562     0.4426  -3.742
speciesPESU          -1.5507     0.3339  -4.644
speciesMYLU          -0.9636     0.1980  -4.866
speciesEPFU          -2.1173     0.3475  -6.092
twinter              -1.4356     0.5450  -2.634
speciesPESU:twinter   1.0992     0.4214   2.609
speciesMYLU:twinter   0.7705     0.2815   2.737
speciesEPFU:twinter   0.5327     0.4265   1.249

Correlation of Fixed Effects:
            (Intr) spPESU spMYLU spEPFU twintr sPESU: sMYLU:
speciesPESU -0.224                                          
speciesMYLU -0.274  0.387                                   
speciesEPFU -0.133  0.153  0.305                            
twinter     -0.812  0.182  0.222  0.108                     
spcsPESU:tw  0.177 -0.792 -0.307 -0.121 -0.271              
spcsMYLU:tw  0.193 -0.272 -0.703 -0.215 -0.328  0.422       
spcsEPFU:tw  0.108 -0.125 -0.249 -0.815 -0.218  0.233  0.374
```


```
emmeans(mod3, pairwise ~ t|species)
```


```
$emmeans
species = MYSE:
 t      emmean    SE   df lower.CL upper.CL
 fall    -1.66 0.443 20.2    -2.58   -0.734
 winter  -3.09 0.318 32.3    -3.74   -2.443

species = PESU:
 t      emmean    SE   df lower.CL upper.CL
 fall    -3.21 0.492 29.8    -4.21   -2.202
 winter  -3.54 0.330 36.5    -4.21   -2.873

species = MYLU:
 t      emmean    SE   df lower.CL upper.CL
 fall    -2.62 0.433 18.5    -3.53   -1.713
 winter  -3.28 0.298 24.9    -3.90   -2.671

species = EPFU:
 t      emmean    SE   df lower.CL upper.CL
 fall    -3.77 0.525 39.5    -4.84   -2.711
 winter  -4.68 0.319 31.6    -5.33   -4.025

Degrees-of-freedom method: kenward-roger 
Confidence level used: 0.95 

$contrasts
species = MYSE:
 contrast      estimate    SE   df t.ratio p.value
 fall - winter    1.436 0.545 23.5 2.632   0.0147 

species = PESU:
 contrast      estimate    SE   df t.ratio p.value
 fall - winter    0.336 0.593 31.7 0.568   0.5743 

species = MYLU:
 contrast      estimate    SE   df t.ratio p.value
 fall - winter    0.665 0.525 20.2 1.266   0.2197 

species = EPFU:
 contrast      estimate    SE   df t.ratio p.value
 fall - winter    0.903 0.615 37.1 1.468   0.1504 

Degrees-of-freedom method: kenward-roger
```

#### 3.3 lambda in late winter


```
mod4=lmer(log.lambda~species*t + (1|site), data=data.trim.late.lambda);summary(mod4)
```


```
Linear mixed model fit by REML ['lmerMod']
Formula: log.lambda ~ species * t + (1 | site)
   Data: data.trim.late.lambda

REML criterion at convergence: 31.9

Scaled residuals: 
     Min       1Q   Median       3Q      Max 
-1.80892 -0.55960 -0.04786  0.60157  1.81107 

Random effects:
 Groups   Name        Variance Std.Dev.
 site     (Intercept) 0.05463  0.2337  
 Residual             0.04941  0.2223  
Number of obs: 60, groups:  site, 20

Fixed effects:
                    Estimate Std. Error t value
(Intercept)          -0.2439     0.1317  -1.852
speciesEPFU           0.5945     0.1645   3.614
speciesMYLU           0.3860     0.1283   3.008
speciesPESU           0.1889     0.1477   1.279
twinter               0.4350     0.1655   2.628
speciesEPFU:twinter  -0.5674     0.1970  -2.880
speciesMYLU:twinter  -0.4112     0.1636  -2.513
speciesPESU:twinter  -0.2431     0.1818  -1.337

Correlation of Fixed Effects:
            (Intr) spEPFU spMYLU spPESU twintr sEPFU: sMYLU:
speciesEPFU -0.380                                          
speciesMYLU -0.487  0.390                                   
speciesPESU -0.423  0.340  0.434                            
twinter     -0.796  0.303  0.388  0.337                     
spcsEPFU:tw  0.317 -0.835 -0.326 -0.284 -0.445              
spcsMYLU:tw  0.382 -0.306 -0.784 -0.341 -0.528  0.443       
spcsPESU:tw  0.344 -0.276 -0.353 -0.812 -0.472  0.394  0.478
```


`r`r emmeans(mod4, pairwise~t|species)


```
$emmeans
species = MYSE:
 t       emmean     SE   df lower.CL upper.CL
 fall   -0.3630 0.1460 31.4  -0.6605  -0.0654
 winter  0.1644 0.1140 40.0  -0.0660   0.3949

species = EPFU:
 t       emmean     SE   df lower.CL upper.CL
 fall    0.2625 0.2005 42.0  -0.1422   0.6672
 winter  0.2426 0.0999 34.6   0.0397   0.4455

species = MYLU:
 t       emmean     SE   df lower.CL upper.CL
 fall    0.0319 0.1460 31.4  -0.2656   0.3294
 winter  0.1758 0.1048 36.2  -0.0367   0.3884

species = PESU:
 t       emmean     SE   df lower.CL upper.CL
 fall   -0.0397 0.1565 35.9  -0.3571   0.2778
 winter  0.1123 0.1091 38.2  -0.1085   0.3331

Degrees-of-freedom method: kenward-roger 
Confidence level used: 0.95 

$contrasts
species = MYSE:
 contrast      estimate    SE   df t.ratio p.value
 fall - winter  -0.5274 0.185 35.0 -2.848  0.0073 

species = EPFU:
 contrast      estimate    SE   df t.ratio p.value
 fall - winter   0.0199 0.224 41.8  0.089  0.9296 

species = MYLU:
 contrast      estimate    SE   df t.ratio p.value
 fall - winter  -0.1439 0.180 33.1 -0.801  0.4288 

species = PESU:
 contrast      estimate    SE   df t.ratio p.value
 fall - winter  -0.1519 0.191 36.7 -0.796  0.4309 

Degrees-of-freedom method: kenward-roger
```

### 4 CONTINUOUS ANALYSIS - SAME AS ABOVE BUT CONTINUOUS

#### 4.1 how are late winter loads affected by early prevalence?


```
mod6=lmer(lgdL~species+early.prev +(1|site), data=mw0.trim.late);summary(mod6)
```


```
Linear mixed model fit by REML ['lmerMod']
Formula: lgdL ~ species + early.prev + (1 | site)
   Data: mw0.trim.late

REML criterion at convergence: 1261.6

Scaled residuals: 
     Min       1Q   Median       3Q      Max 
-2.93185 -0.67037  0.01515  0.65539  2.98943 

Random effects:
 Groups   Name        Variance Std.Dev.
 site     (Intercept) 1.049    1.024   
 Residual             1.262    1.123   
Number of obs: 394, groups:  site, 19

Fixed effects:
            Estimate Std. Error t value
(Intercept)  -2.7819     0.2877  -9.668
speciesPESU  -0.6462     0.2339  -2.762
speciesMYLU  -0.2910     0.1773  -1.641
speciesEPFU  -1.9322     0.2700  -7.155
early.prev    0.8494     0.2893   2.936

Correlation of Fixed Effects:
            (Intr) spPESU spMYLU spEPFU
speciesPESU -0.401                     
speciesMYLU -0.459  0.520              
speciesEPFU -0.341  0.379  0.438       
early.prev  -0.335  0.393  0.558  0.341
```

#### 4.2 how is late winter prevalence affected by early prevalence?


```
mod5=glmer(gd~species+early.prev +(1|site),weights=N, data=data.trim.late,family="binomial");summary(mod5)
```


```
Generalized linear mixed model fit by maximum likelihood (Laplace Approximation) ['glmerMod']
 Family: binomial  ( logit )
Formula: gd ~ species + early.prev + (1 | site)
   Data: data.trim.late
Weights: N

     AIC      BIC   logLik deviance df.resid 
   204.3    216.3    -96.1    192.3       49 

Scaled residuals: 
    Min      1Q  Median      3Q     Max 
-1.9955 -0.5332  0.1541  0.6788  1.6161 

Random effects:
 Groups Name        Variance Std.Dev.
 site   (Intercept) 4.231    2.057   
Number of obs: 55, groups:  site, 20

Fixed effects:
            Estimate Std. Error z value Pr(>|z|)    
(Intercept)   1.6992     0.6009   2.828  0.00469 ** 
speciesEPFU  -2.7521     0.6720  -4.095 4.21e-05 ***
speciesMYLU  -0.2661     0.3614  -0.736  0.46151    
speciesPESU  -2.5151     0.4114  -6.114 9.71e-10 ***
early.prev   12.4435     6.2987   1.976  0.04820 *  
---
Signif. codes:  0 ‘***’ 0.001 ‘**’ 0.01 ‘*’ 0.05 ‘.’ 0.1 ‘ ’ 1

Correlation of Fixed Effects:
            (Intr) spEPFU spMYLU spPESU
speciesEPFU -0.388                     
speciesMYLU -0.464  0.425              
speciesPESU -0.492  0.445  0.686       
early.prev  -0.288  0.123  0.144  0.220
```

#### 4.3 how are late winter impacts affected by early prevalence?


```
mod6=lmer(log.lambda~species+early.prev +(1|site), data=data.trim.late.lambda);summary(mod6)
```


```
Linear mixed model fit by REML ['lmerMod']
Formula: log.lambda ~ species + early.prev + (1 | site)
   Data: data.trim.late.lambda

REML criterion at convergence: 24.1

Scaled residuals: 
     Min       1Q   Median       3Q      Max 
-2.31360 -0.38344 -0.04979  0.62521  2.23076 

Random effects:
 Groups   Name        Variance Std.Dev.
 site     (Intercept) 0.05378  0.2319  
 Residual             0.05162  0.2272  
Number of obs: 49, groups:  site, 18

Fixed effects:
            Estimate Std. Error t value
(Intercept) -0.03862    0.08458  -0.457
speciesEPFU  0.20691    0.11698   1.769
speciesMYLU  0.13551    0.08910   1.521
speciesPESU  0.08589    0.09140   0.940
early.prev   0.48082    0.70962   0.678

Correlation of Fixed Effects:
            (Intr) spEPFU spMYLU spPESU
speciesEPFU -0.433                     
speciesMYLU -0.482  0.398              
speciesPESU -0.516  0.418  0.483       
early.prev  -0.158 -0.045 -0.234  0.006
```

### 5 OVERWINTER MOBILITY

#### 5.1 Changes in counts over winter - site


```
summary(mod.overwinter.movements2)
```


```
Linear mixed model fit by REML ['lmerMod']
Formula: log.overwinter.lambda ~ log.early.winter.count + species + (1 |      site)
   Data: preWNS

REML criterion at convergence: 46.4

Scaled residuals: 
     Min       1Q   Median       3Q      Max 
-2.83773 -0.48121  0.02284  0.56248  2.99240 

Random effects:
 Groups   Name        Variance Std.Dev.
 site     (Intercept) 0.02909  0.1706  
 Residual             0.07031  0.2652  
Number of obs: 88, groups:  site, 16

Fixed effects:
                        Estimate Std. Error t value
(Intercept)             0.277126   0.113676   2.438
log.early.winter.count -0.152472   0.057270  -2.662
speciesMYLU             0.018150   0.090899   0.200
speciesMYSE            -0.103511   0.100143  -1.034
speciesPESU            -0.009488   0.088944  -0.107

Correlation of Fixed Effects:
            (Intr) lg.r.. spMYLU spMYSE
lg.rly.wnt. -0.744                     
speciesMYLU -0.409  0.004              
speciesMYSE -0.556  0.283  0.521       
speciesPESU -0.571  0.208  0.602  0.568
```

#### 5.2 Does the log transformation of lambda help normality?


```
hist(preWNS$log.overwinter.lambda)
```

#### 5.3 Do model diagnostic plots look reasonable?


```
plot(mod.overwinter.movements2)
```

#### 5.4 If we preserve structure of data and use Gamma distribution do we get a similar finding?


```
summary(mod.overwinter.movements2g)
```


```
Generalized linear mixed model fit by maximum likelihood (Laplace Approximation) ['glmerMod']
 Family: Gamma  ( log )
Formula: overwinter.lambda ~ log.early.winter.count + species + (1 | site)
   Data: preWNS

     AIC      BIC   logLik deviance df.resid 
   200.0    217.4    -93.0    186.0       81 

Scaled residuals: 
    Min      1Q  Median      3Q     Max 
-1.4921 -0.5895 -0.0835  0.4068  3.9362 

Random effects:
 Groups   Name        Variance Std.Dev.
 site     (Intercept) 0.2064   0.4544  
 Residual             0.3322   0.5763  
Number of obs: 88, groups:  site, 16

Fixed effects:
                       Estimate Std. Error t value Pr(>|z|)   
(Intercept)             0.91237    0.28271   3.227  0.00125 **
log.early.winter.count -0.38356    0.12130  -3.162  0.00157 **
speciesMYLU            -0.05143    0.19464  -0.264  0.79160   
speciesMYSE            -0.30116    0.22053  -1.366  0.17207   
speciesPESU            -0.14827    0.19353  -0.766  0.44360   
---
Signif. codes:  0 ‘***’ 0.001 ‘**’ 0.01 ‘*’ 0.05 ‘.’ 0.1 ‘ ’ 1

Correlation of Fixed Effects:
            (Intr) lg.r.. spMYLU spMYSE
lg.rly.wnt. -0.654                     
speciesMYLU -0.409  0.070              
speciesMYSE -0.520  0.322  0.580       
speciesPESU -0.537  0.254  0.652  0.637
```

### 6 NUMEROUS COVARIATES AND THE EFFECT ON TIMING OF DETECTION

#### 6.1 temperature


```
summary(gob8)
```


```
Call:
glm(formula = pd.arrival ~ mean.temp, family = "binomial", data = site.dat)

Deviance Residuals: 
    Min       1Q   Median       3Q      Max  
-1.1451  -0.9399  -0.7092   1.4147   1.6224  

Coefficients:
            Estimate Std. Error z value Pr(>|z|)
(Intercept)  -2.1649     1.7413  -1.243    0.214
mean.temp     0.2083     0.2313   0.900    0.368

(Dispersion parameter for binomial family taken to be 1)

    Null deviance: 26.734  on 20  degrees of freedom
Residual deviance: 25.839  on 19  degrees of freedom
  (1 observation deleted due to missingness)
AIC: 29.839

Number of Fisher Scoring iterations: 4
```

#### 6.2 vapor pressure deficit


```
summary(gob9a)
```


```
Call:
glm(formula = pd.arrival ~ mean.vpd, family = "binomial", data = site.dat)

Deviance Residuals: 
    Min       1Q   Median       3Q      Max  
-1.1390  -0.9353  -0.7392   1.2213   1.7577  

Coefficients:
             Estimate Std. Error z value Pr(>|z|)
(Intercept)  -0.04919    0.68436  -0.072    0.943
mean.vpd    -17.61464   15.46895  -1.139    0.255

(Dispersion parameter for binomial family taken to be 1)

    Null deviance: 26.734  on 20  degrees of freedom
Residual deviance: 25.079  on 19  degrees of freedom
  (1 observation deleted due to missingness)
AIC: 29.079

Number of Fisher Scoring iterations: 4
```

#### 6.3 total bats


```
summary(gob1)
```


```
Call:
glm(formula = pd.arrival ~ log10(total), family = "binomial", 
    data = site.dat)

Deviance Residuals: 
    Min       1Q   Median       3Q      Max  
-1.0612  -0.8776  -0.8166   1.4364   1.5859  

Coefficients:
             Estimate Std. Error z value Pr(>|z|)
(Intercept)   -1.2752     1.3300  -0.959    0.338
log10(total)   0.2384     0.5736   0.416    0.678

(Dispersion parameter for binomial family taken to be 1)

    Null deviance: 27.522  on 21  degrees of freedom
Residual deviance: 27.350  on 20  degrees of freedom
AIC: 31.35

Number of Fisher Scoring iterations: 4
```

#### 6.4 number of MYLU


```
summary(gob4)
```


```
Call:
glm(formula = pd.arrival ~ sum.mylu, family = "binomial", data = site.dat)

Deviance Residuals: 
    Min       1Q   Median       3Q      Max  
-0.8931  -0.8930  -0.8903   1.4917   1.5964  

Coefficients:
              Estimate Std. Error z value Pr(>|z|)
(Intercept) -7.133e-01  4.785e-01  -1.491    0.136
sum.mylu    -6.548e-05  2.087e-04  -0.314    0.754

(Dispersion parameter for binomial family taken to be 1)

    Null deviance: 27.522  on 21  degrees of freedom
Residual deviance: 27.409  on 20  degrees of freedom
AIC: 31.409

Number of Fisher Scoring iterations: 4
```

#### 6.5 number of EPFU


```
summary(gob6)
```


```
Call:
glm(formula = pd.arrival ~ sum.epfu, family = "binomial", data = site.dat)

Deviance Residuals: 
    Min       1Q   Median       3Q      Max  
-0.8916  -0.8908  -0.8832   1.4931   1.6046  

Coefficients:
              Estimate Std. Error z value Pr(>|z|)
(Intercept) -0.7172423  0.4970035  -1.443    0.149
sum.epfu    -0.0008242  0.0037308  -0.221    0.825

(Dispersion parameter for binomial family taken to be 1)

    Null deviance: 27.522  on 21  degrees of freedom
Residual deviance: 27.470  on 20  degrees of freedom
AIC: 31.47

Number of Fisher Scoring iterations: 4
```

#### 6.6 overwinter mobility


```
summary(gob14)
```


```
Call:
glm(formula = pd.arrival ~ log.overwinter.lambda, family = "binomial", 
    data = site.dat)

Deviance Residuals: 
    Min       1Q   Median       3Q      Max  
-1.0806  -0.7973  -0.7248   0.1033   1.6431  

Coefficients:
                      Estimate Std. Error z value Pr(>|z|)
(Intercept)            -0.8887     0.6109  -1.455    0.146
log.overwinter.lambda  -2.1356     2.6492  -0.806    0.420

(Dispersion parameter for binomial family taken to be 1)

    Null deviance: 17.995  on 15  degrees of freedom
Residual deviance: 17.161  on 14  degrees of freedom
  (6 observations deleted due to missingness)
AIC: 21.161

Number of Fisher Scoring iterations: 4
```

#### 6.7 effect of early winter sampling date


```
gob15 = glm(pd.arrival~early.pdates, family = "binomial", data=site.dat);summary(gob15)
```


```
Call:
glm(formula = pd.arrival ~ early.pdates, family = "binomial", 
    data = site.dat)

Deviance Residuals: 
    Min       1Q   Median       3Q      Max  
-1.1193  -0.9260  -0.8429   1.3045   1.5832  

Coefficients:
             Estimate Std. Error z value Pr(>|z|)
(Intercept)   -9.9006    17.3967  -0.569    0.569
early.pdates   0.8049     1.5068   0.534    0.593

(Dispersion parameter for binomial family taken to be 1)

    Null deviance: 25.898  on 19  degrees of freedom
Residual deviance: 25.613  on 18  degrees of freedom
  (2 observations deleted due to missingness)
AIC: 29.613

Number of Fisher Scoring iterations: 4
```

#### 6.8 species richness


```
gob16 = glm(pd.arrival~spec.num, family = "binomial", data=site.dat);summary(gob16)
```


```
Call:
glm(formula = pd.arrival ~ spec.num, family = "binomial", data = site.dat)

Deviance Residuals: 
    Min       1Q   Median       3Q      Max  
-1.0358  -0.8487  -0.8487   1.3259   1.7686  

Coefficients:
            Estimate Std. Error z value Pr(>|z|)
(Intercept)  -2.3155     2.0912  -1.107    0.268
spec.num      0.4932     0.6356   0.776    0.438

(Dispersion parameter for binomial family taken to be 1)

    Null deviance: 27.522  on 21  degrees of freedom
Residual deviance: 26.871  on 20  degrees of freedom
AIC: 30.871

Number of Fisher Scoring iterations: 4
```
